# T cell AIM2 Exacerbates Lung Inflammation in OVA-LPS and HDM-induced Asthma Models

**DOI:** 10.1101/2025.07.02.662837

**Authors:** Wei-Chun Chou, Martin Hsu, Corey M. Jania, Elizabeth Holley-Guthrie, John A. Wrobel, Kaixin Liang, Alessandra Livraghi-Butrico, Stephen L. Tilley, Yisong Y. Wan, Jenny P.-Y. Ting

## Abstract

Asthma is a complex airway inflammatory disease characterized by immune dysregulation, with diverse cellular contributors to inflammation that result in airway hyperresponsiveness (AHR) and excessive immune responses. The role of the inflammasome in asthma is conflicted. AIM2 is an innate immune receptor that activates the inflammasome through DNA binding. The clinical relevance of AIM2 in asthma is supported by analysis of blood or endobronchial biopsies from patients with severe asthma, showing increased AIM2 but not NLRP3 expression. We investigated the role of AIM2 in allergic asthma using ovalbumin (OVA)-LPS and house dust mite (HDM)- induced models. AIM2 expression was elevated in lung homogenates and bronchoalveolar lavage fluid (BALF) cells from allergen-induced groups compared to controls. In the OVA-LPS model, deletion of *Aim2* led to reduced airway smooth muscle actin expression and DNA damage, contributing to decreased AHR and lung inflammation in whole body *Aim2^−/−^* mice. Surprisingly, cell-specific deletion in CD4^+^ T cells but not regulatory T cells or myeloid cells significantly affected AHR, indicating AIM2 in CD4^+^ T cells is a main driver of pathogenesis. Additionally, in HDM-induced asthma, *Aim2^−/−^* mice also exhibited reduced AHR. Cell-specific deletion reveals that AIM2 in CD4^+^ T cells promotes AHR and cytokine production, while AIM2 in myeloid cells modulates IgG1 levels and IL-13-producing CD4^+^ T cells, and the percentage of alveolar macrophages in BALF. These findings reveal that AIM2 differentially regulates HDM-induced allergic asthma through distinct exacerbating roles in T cells and myeloid cells, highlighting its potential as a therapeutic target.

**Significance:** Allergic asthma is a complex, heterogeneous disease characterized by airway inflammation and hyper-responsiveness. AIM2, an inflammasome activated by DNA, has emerged as a critical regulator of inflammatory diseases. In humans, AIM2 expression is elevated in severe asthma. *In vivo* studies highlight AIM2 as a key modulator of allergic asthma in both OVA-LPS and HDM-induced models. Surprisingly, its functional relevance in both models is mediated by its intrinsic role in T cells, while a modest role in myeloid cells is restricted to the HDM model. AIM2 promotes airway hyper-responsiveness and lung inflammation through enhancing type II cytokines, immune subpopulations and DNA damage response. These findings suggest AIM2 as a biomarker and therapeutic target for managing asthma, particularly in severe asthma subtypes.

## Introduction

Asthma is a chronic inflammatory respiratory disease caused by complex interactions between innate and adaptive immune cells, leading to airway inflammation, hyperresponsiveness (AHR), and remodeling. Upon allergen exposure, epithelial cells release cytokines such as IL-25, IL-33, and TSLP, which activate dendritic cells, innate lymphoid cells (ILCs), and other innate immune cells. These signaling pathways drive a Th2-dominant adaptive immune response, characterized by CD4^+^ Th2 cells producing IL-4, IL-5, and IL-13. This response leads to eosinophil recruitment, IgE production, and increased mucus secretion(*1–5*). Additionally, the Th2 endotypes of Th2-high or Th2-low/ non-Th2 pathways, including Th17 cells and neutrophilic inflammation, contribute to the heterogeneity of asthma(*2, 3, 5*). Although asthma has been studied extensively by using different allergen-induced mouse models(*6–11*), the underlying mechanisms driving its etiology and pathogenesis of the disease remain unclear.

Previous studies have highlighted the involvement of inflammasome activation, a multi-protein complex consisting of a ligand sensor, ASC (apoptosis-associated speck-like protein containing a caspase recruitment domain) and caspase-1, in triggering inflammatory responses, particularly by processing and secreting pro-inflammatory cytokines such as IL-1β and IL-18, under various asthmatic conditions (*12–16*). Among the various inflammasomes, the NLRP3 (NOD-like receptor family pyrin domain-containing 3) inflammasome has been extensively studied in the context of asthma, demonstrating its role in both inflammasome-dependent and independent mechanisms across different cell types in various experimental mouse models of asthma(*17–26*). For instance, while an initial report indicates the importance of NLRP3 in alum adjuvant induced lung allergy(*27*), we and others did not observe this effect(*20, 28, 29*). However, in other asthma models the role of NLRP3 has been observed, suggesting that NLRP3 may be important only in certain asthma or airway inflammation models(*30, 31*). Additionally, reports have shown that the inhibition of IL-1β and caspase-1, as well as treatment with NLRP3-specific inhibitors such as MCC950, dapansutrile (OLT1177), and RRx-001, suppresses airway inflammation in OVA- or HDM-induced allergic asthma models through inflammasome-dependent mechanisms (*24, 25, 32*). In contrast, a report indicated that OVA challenge alone fails to increase IL-1β and caspase-1 expression, although sevoflurane, similar to MCC950, ameliorated allergic airway inflammation by inhibiting Th2 responses and NLRP3 expression (*29*).

Despite the well-documented role of NLRP3 in asthma, the potential involvement of other inflammasomes in the disease has not been fully explored. AIM2 (Absent in Melanoma 2) is another inflammasome sensor that recognizes cytosolic double-stranded DNA (dsDNA) and activates the inflammasome, leading to the production of IL-1β and IL-18. AIM2 has been extensively studied in various cell types within the contexts of microbial infections, cancer, and autoimmune diseases, where it functions through both inflammasome-dependent and independent mechanisms (*33–43*). Additionally, in lung inflammation, AIM2 has been associated with influenza and mycobacterium tuberculosis infection (*44, 45*), chronic obstructive pulmonary disease (COPD)(*46*), pulmonary fibrosis (*47*). More recently, it has also been implicated in contributing to epithelial dysfunction in infant asthma (*48*). However, its role in asthma pathogenesis is less well-defined.

In this study, we tested the hypothesis that AIM2 contributes to lung inflammation by functioning in immune cells in asthma. Specifically, we investigated the role of AIM2 in asthma by examining its expression and function in two distinct murine models of asthma: OVA-LPS-induced and HDM-induced models. These models mimic key aspects of human heterogenous endotypes of asthma, including eosinophilic and neutrophilic inflammation, and airway hyperresponsiveness (AHR). By utilizing cell-specific deletion models, we found that AIM2 functions in a T cell-intrinsic or myeloid cell-intrinsic manner to modulate asthma severity. Our findings provide insight into the cell-specific contributions of AIM2 in allergic asthma and suggest that AIM2 may be a critical driver of lung inflammation in certain asthma phenotypes. Moreover, the differential expression of AIM2 observed in human severe asthma and neutrophilic versus paucigranulocytic asthma samples highlights its potential role as a biomarker for asthma. Together, these results establish AIM2 as a potential therapeutic target for modulating immune responses in allergic asthma.

## Results

### Heightened AIM2 Expression in Human Samples from Asthmatic Patients

Increased *AIM2* expression was previously observed in severe asthma pediatric patients compared to normal controls according to the gene expression profiles analyzed by a recent report using published database GSE27011 in severe therapy-resistant childhood asthma(*48*). To explore whether *AIM2* has differential expression level in different human asthmatic subtypes, we further analyzed two public database GSE69683(*49*) and GSE143303 (*50*). Our analysis of publicly available transcriptomic datasets shows that *AIM2* expression is significantly elevated in both the peripheral blood of patients with severe asthma (log₂ fold change 0.207; adjusted p = 0.022; GSE69683) (Fig. 1A-C) and the bronchial tissue of patients with neutrophilic asthma (log₂ fold change 0.571; adjusted p = 0.017; GSE143303), compared to respective controls (Fig. 1D-F). By contrast, *NLRP3* expression did not differ significantly between groups. These findings suggest that *AIM2*, rather than *NLRP3*, may serve as a key molecular marker associated with severe and neutrophilic asthma phenotypes.

**Fig. 1.**
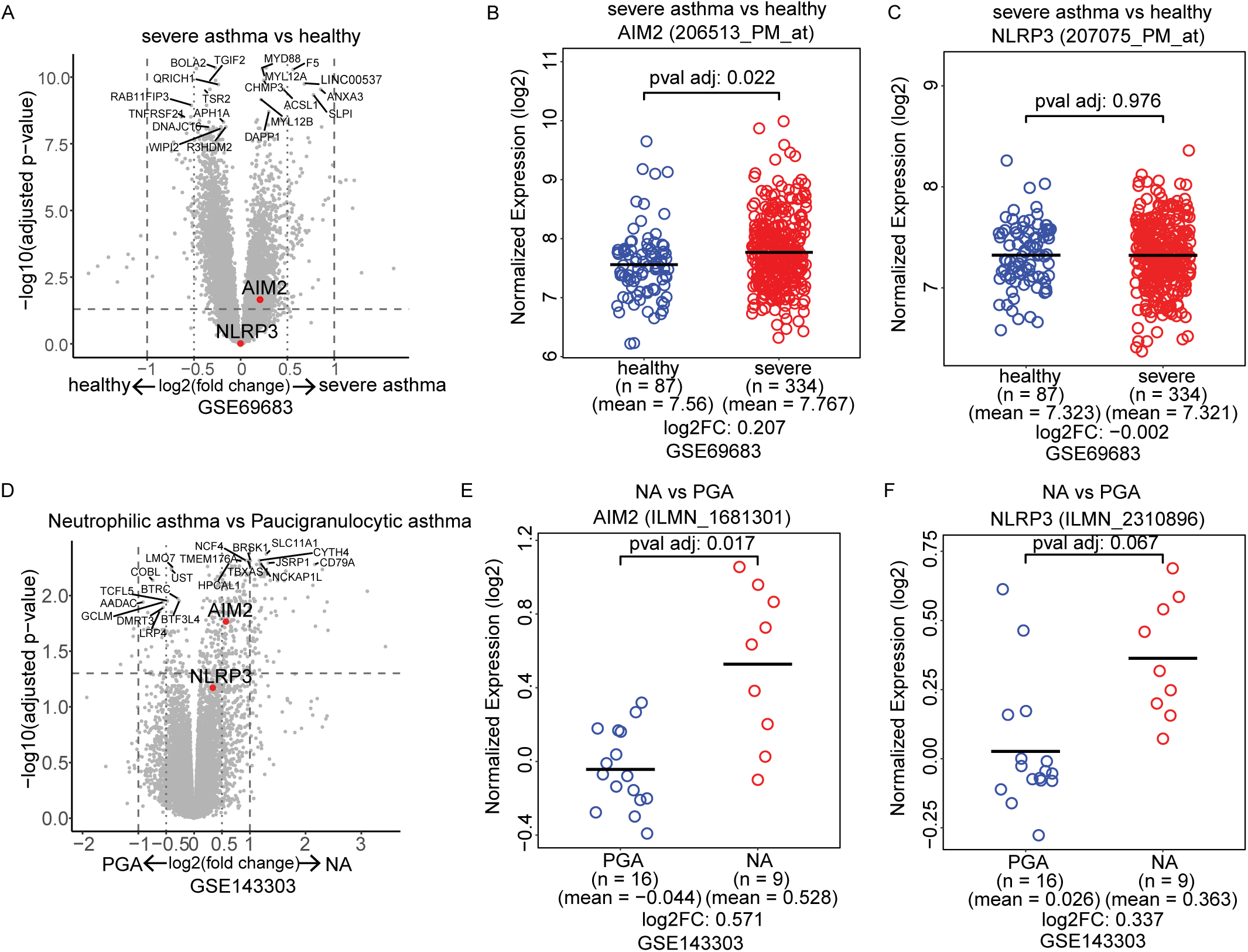
*AIM2* gene expression in human asthma samples. (A) Volcano plot showing differential expression gene expression between blood samples from patients with severe asthma (n = 334) and individuals without asthma (n = 87) from the publicly available microarray dataset (GSE69683). *AIM2* and *NLRP3* are indicated with red dots. The x-axis represents log2 fold change (severe asthma / healthy) and the y-axis represents the adjusted p-value. Sample level gene expression comparison for (B) *AIM2* and (C) *NLRP3*. In each plot, the y-axis indicates the normalized log2 expression values for each sample group on the x-axis, with each circle representing an individual sample. The severe asthma samples are colored as red circles and the healthy samples are colored as blue samples. A black bar indicates the mean expression of each sample group. The log2 fold change (severe asthma / healthy) and adjusted p-value are listed in each plot. (D) Volcano plot showing differential expression gene expression between endobronchial biopsy samples from adults with neutrophilic asthma (NA) (n = 9) and paucigranulocytic asthma (PGA) (n = 16) from the publicly available microarray dataset (GSE143303). *AIM2* and *NLRP3* are indicated with red dots. The x-axis represents log2 fold change (severe asthma / healthy) and the y-axis represents the adjusted p-value. Sample level gene expression comparison for (E) *AIM2* and (F) *NLRP3*. In each plot, the y-axis indicates the normalized log2 expression values for each sample group on the x-axis, with each circle representing an individual sample. The NA samples are colored as red circles and the PGA samples are colored as blue circles. A black bar indicates the mean expression of each sample group. The log2 fold change (NA/PGA) and adjusted p-value are listed in each plot.

### AIM2 Exhibits Increased Expression and Exacerbates Lung Inflammation in the OVA-LPS Model

To investigate the functional role of AIM2 in allergic asthma, we first utilized the ovalbumin (OVA)-LPS-induced allergic asthma murine model (Fig. 2A). *Aim2* expression was elevated in both total lung homogenates and bronchoalveolar lavage fluid (BALF) cells from the OVA-LPS-induced group compared to the PBS control group (Fig. 2B). OVA-LPS stimulation significantly increased airway hyperresponsiveness (AHR) measured by different parameters including total airway resistance (Rrs), upper proximal airway resistance (Rn) and lower distal airway resistance (G) in WT mice upon dose-dependent methacholine challenge, while modestly reduced AHR was observed in *Aim2^−/−^* mice (Fig. 2C). Chemokine (C-C motif) ligand 2 (CCL2) level is reported to be increased in primary <u>a</u>irway <u>s</u>mooth <u>m</u>uscle (ASM) supernatants from asthmatics compared with healthy controls (*51*). Interestingly, *ccl2* expression was reduced in *Aim2^−/−^* lung lysates, however, Th1/Th2/Th17 cytokines (*Ifng, Il4, Il5, Il13, Il17a*), inflammasome genes (*Il1b and Il18*) and others were not altered at the mRNA level (Fig. S1A). Additionally, IL-4, CCL2, and TNF production were significantly reduced at the protein level (Fig. 2D), while other cytokines (IL-5, IP-10, IL-1β, IL-18, and eotaxin) (Fig. S1B) as well as the ratio of IgG1/IgG2a (Fig. S1C) were not changed in *Aim2^−/−^* BALF compared to WT BALF. As OVA-LPS model is known as a mixed Th1/Th2/Th17 allergic asthma model (*3*), we next assessed which cytokine-producing T cell profile was involved in the activation of AIM2 in the BALF in the OVA-LPS model. We found reduced CD4^+^Foxp3^+^ regulatory T cells (Fig. 2E), increased IFNγ-producing- (Fig. 2F) but reduced IL-13-producing-CD4^+^ T cells (Fig. 2G) in *Aim2^−/−^* BALF. However, there is no difference in IL-17-producing-CD4^+^ T cells (Fig. 2G). There is also no difference in CD8^+^ IFNγ-producing-, IL-13-producing-, or IL-17-producing T cells (Fig. 2H and I) between WT and *Aim2^−/−^* BALF, suggesting AIM2 is not affecting CD8 cells in OVA-LPS model.

**Fig. 2.**
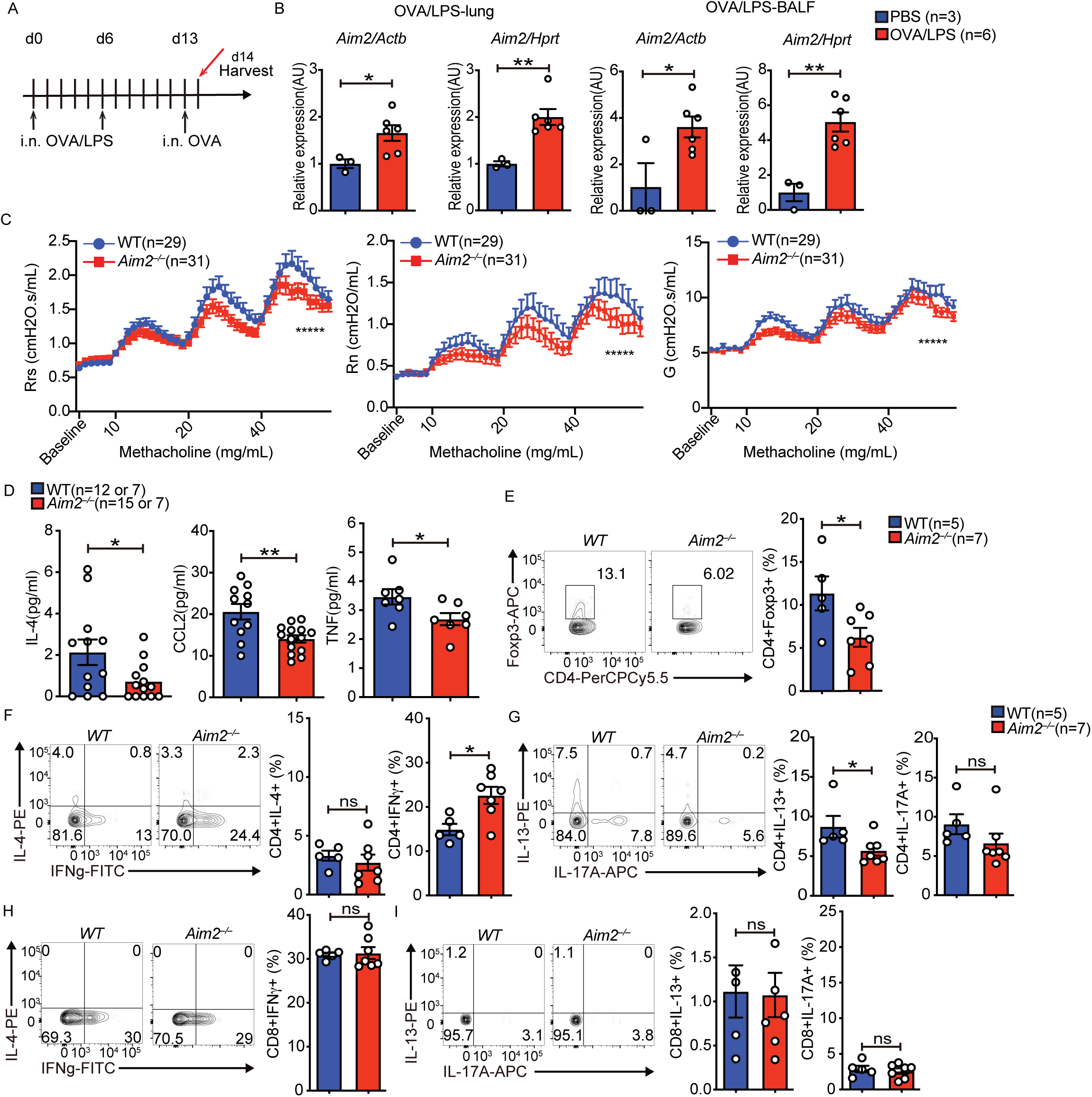
AIM2 exhibits increased expression and exacerbates lung inflammation in the OVA-LPS model. *WT* and *Aim2^−/−^* mice were intranasally sensitized to OVA (100 μg) using LPS (100 ng) as the adjuvant, and then intranasally challenged with a single dose of OVA (100 μg) on the day before harvest. (A) Schematic of allergic sensitizations, challenge and harvest. (B) *Aim2* expression in the lung (left panel) and in the bronchoalveolar lavage fluid (BALF) (right panel). (C) Airway hyperresponsiveness (AHR) was measured upon administration of indicated methacholine doses, and measured by three parameters, including resistance of total airway (Rrs), proximal, large airway (Rn), and distal small airway (G) in mice. *WT* n = 29, *Aim2^−/−^* n = 31. (D) Protein concentration of different cytokines in the BALF was measured by ELISA. (E to I) Flow cytometry analysis of Foxp3 Treg cells (E), IL-4, IFNγ, IL-13, and IL-17-secreting CD4^+^ (F and G) or CD8^+^ (H and I) T cells. Data are representative of three independent experiments. Data shown represent mean values ± SEM. *p < 0.05; **p < 0.01; *** p < 0.001; **** p < 0.0001. ns: p >0.05; not significant. Student t-test or Two-way ANOVA with Sidak multiple comparison tests.

### Reduced α-SMA Expression and DNA Damage in AIM2-Deficient Mice in the OVA-LPS Model

Given the findings in OVA-LPS-induced asthmatic mice, which demonstrated a significant reduction in CCL2 levels at both the mRNA and protein levels in *Aim2^−/−^* lungs and BALF, along with a previous report showing elevated of serum and BALF CCL2 levels released by ASM in asthmatic patient (*51*), we investigated whether the absence of AIM2 also affects the expression of airway α-<u>s</u>mooth <u>m</u>uscle <u>a</u>ctin (α-SMA). Immunoblotting of protein samples from lung homogenates revealed a decrease in α-SMA expression in *Aim2^−/−^* lungs (Fig. 3A). Notably, AIM2 was demonstrated to colocalize with γH2A.X (the activated phosphorylation form of H2A.X at Ser139 site) and enhance its expression in response to ionizing radiation-induced DNA damage (*52*). To evaluate whether AIM2 modulates activation of histone H2A.X in OVA-LPS-induced asthma, which was previously reported to be increased in HDM-induced DNA damage in asthmatic mouse and human lung tissues compared to healthy controls (*53*), we examined whether the expression of γH2A.X was altered in the absence of AIM2. γH2A.X expression was decreased in *Aim2^−/−^* lungs upon OVA-LPS stimulation (Fig. 3B). To test if AIM2 functions as an inflammasome in the context of asthma, we sought to determine whether AIM2 influences the activation of caspase-1 and Gasdermin D (GSDMD), both of which are downstream effectors of AIM2 inflammasome activation and contribute to pyroptosis (*54*). However, the activation of caspase-1 and GSDMD was not significantly reduced in the absence of AIM2 as measured by immunoblot and densitometry, although GSDMD showed a reduced trend in *Aim2^-/-^* mice (Fig. 3C). In addition to inflammasome-induced cell death, we further assessed whether AIM2 affects apoptosis in the context of asthma. The activation of both caspase-3 and caspase-7 was also not significantly reduced, which was similarly attributed to decreased total protein levels rather than impaired activation in the absence of AIM2 (Fig. 3D). To further elucidate the underlying mechanisms by which AIM2 may regulate signaling pathways in response to OVA-LPS stimulation, we next examined the activation of STAT1 and STAT3, both of which are implicated in asthma pathogenesis (*55, 56*). Our results showed that the activation of STAT1 and STAT3 was also not significantly diminished, which may be attributed to the reduced total protein levels of each (Fig. 3E). Taken together, our data demonstrated that AIM2 influences the severity of OVA- LPS-induced asthma and is accompanied by increased SMA and DNA damage as revealed by elevated histone H2A.X.

**Fig. 3.**
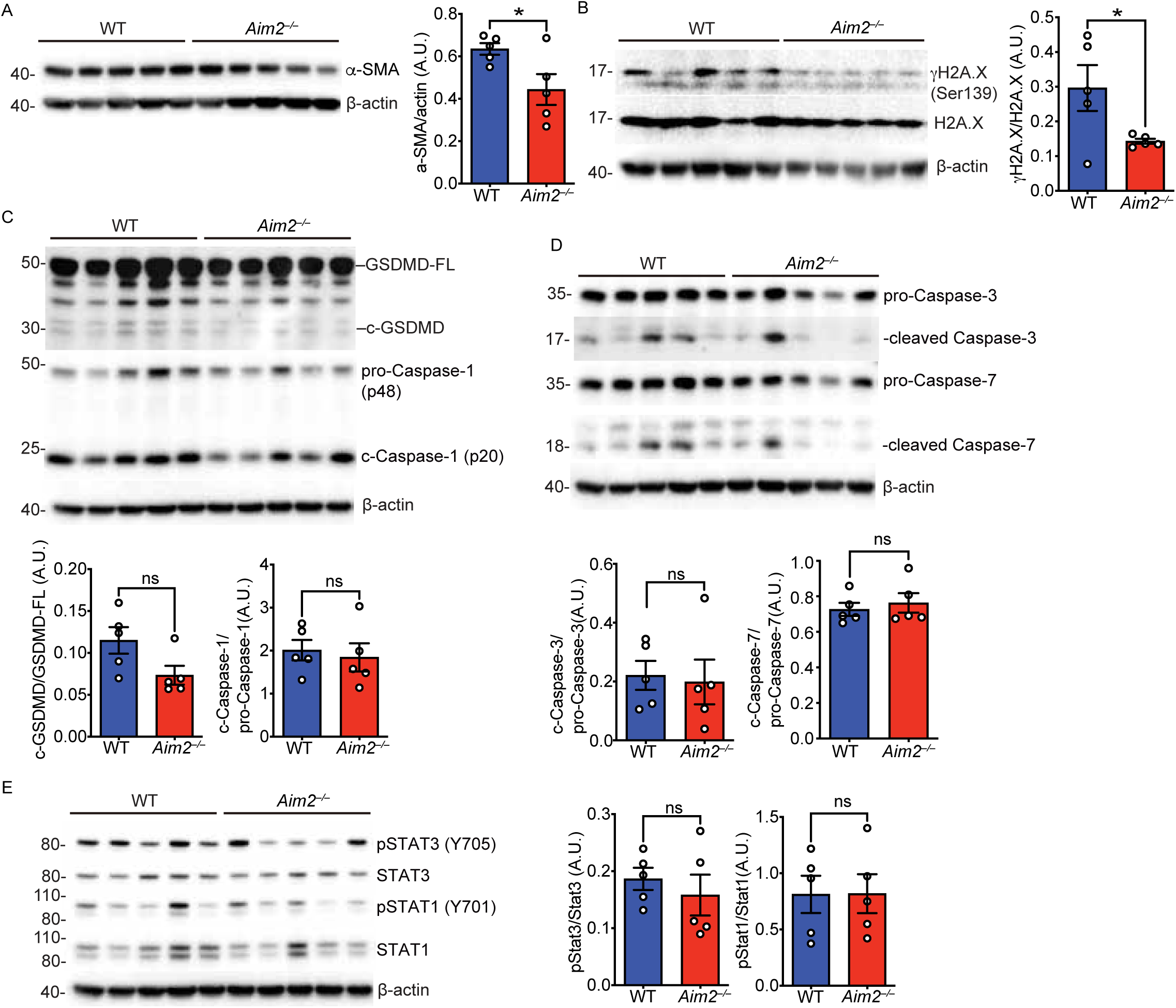
Aim2–/– mice exhibit reduced α-SMA expression, and DNA damage in the OVA-LPS driven lung inflammation model. Immunoblot analysis of different signaling pathways in lung in response to OVA plus LPS, n = 5 per group. (A) α-SMA, (B) phosphorylation of H2A.X., (C) Activation of Gasdermin D (GSDMD) and caspase-1, (D) activation of caspase-3 and caspase-7, and (E) phosphorylation of STAT1 and STAT3.

### T Cell-Intrinsic Deletion of Aim2 Alleviates OVA-LPS-Induced Lung Inflammation

Although AIM2 is primarily described as a myeloid inflammasome receptor, it also has been found to have intrinsic functions in T cells (*43, 57, 58*). We investigated whether AIM2 regulates the severity of OVA-LPS allergic asthma is either CD4 T cell-intrinsic or myeloid cell-intrinsic. To determine the cell-specific role of AIM2 in asthma, we first generated mice with CD4 T cell-specific deletion of *Aim2* by breeding CD4Cre mice with the mouse strain with a floxed *Aim2* allele(*43*). CD4 T cell-specific deletion of *Aim2* was confirmed by western blot (Fig. 4A). We then sensitized and challenged both WT (CD4^Cre^*Aim2^+/+^*) and CD4^Cre^*Aim2*^fl/fl^ by using the same OVA- LPS induced model described in Fig. 2A. CD4^Cre^*Aim2*^fl/fl^ mice displayed dramatically and significantly reduced AHR in response to methacholine challenge compared to CD4^Cre^*Aim2^+/+^*control mice (Fig. 4B). Furthermore, differential gene expression analysis revealed a significant downregulation of asthma-related inflammatory genes (*ccl2*(*59*), *Il17a*(*60*), *Irf4*(*61*), *Pparg*(*62*) and *Il1b*(*63*)) but increased expression of *Il18* in the lung homogenates of CD4^Cre^*Aim2*^fl/fl^ mice (Fig. 4C). Notably, significantly reduced production of IL-4, IL-5, eotaxin, TNF, and IL-6 proteins was observed in CD4^Cre^*Aim2*^fl/fl^ mice (Fig. 4D). The expression of other asthma-related inflammatory genes (Fig. S2A) as well as the level of IL-1β and IL-18 cytokine production (Fig. S2B) were similar between CD4^Cre^*Aim2^+/+^*control mice and CD4^Cre^*Aim2*^fl/fl^mice. There was a trend of reduced *Il13*, *Tnf* and *Cxcl10* transcripts, but these did not reach significance. Additionally, no significant differences were noted in the total production of IgG1, IgG2a, and IgE (Fig. S2C). Following the reduction of type 2 cytokines (IL-4, IL-5, Eotaxin, TNF, and IL-6) in BALF from CD4^Cre^*Aim2*^fl/fl^ mice (Fig. 4D), we next examed whether AIM2 influences Th2 differentiation. CD4⁺ T cells from CD4^Cre^*Aim2*^fl/fl^ mice showed significantly decreased IL-13 production after 4 days of Th2 polarization compared to controls, indicating that CD4^+^ T cell-intrinsic AIM2 promotes Th2 cytokine expression (Fig. 4E). Given that the mTOR-AKT pathway plays a central role in T cell activation, differentiation, and metabolism(*64–66*), and we previously showed AIM2 regulates AKT activation(*37, 43*), we further assessed this pathway in AIM2-deficient T cells. *Aim2*-deficient CD4⁺ T cells exhibited enhanced activation of AKT (S473), while PKCδ/θ (S643/676) phosphorylation was unchanged, suggesting that AIM2 restrains mTOR-AKT signaling during T cell activation, which may contribute to its role in promoting Th2 differentiation (Fig. 4F). Together, these findings indicate that AIM2 in CD4⁺ T cells supports Th2 cytokine production and airway inflammation, at least in part by modulating AKT signaling.

**Fig. 4.**
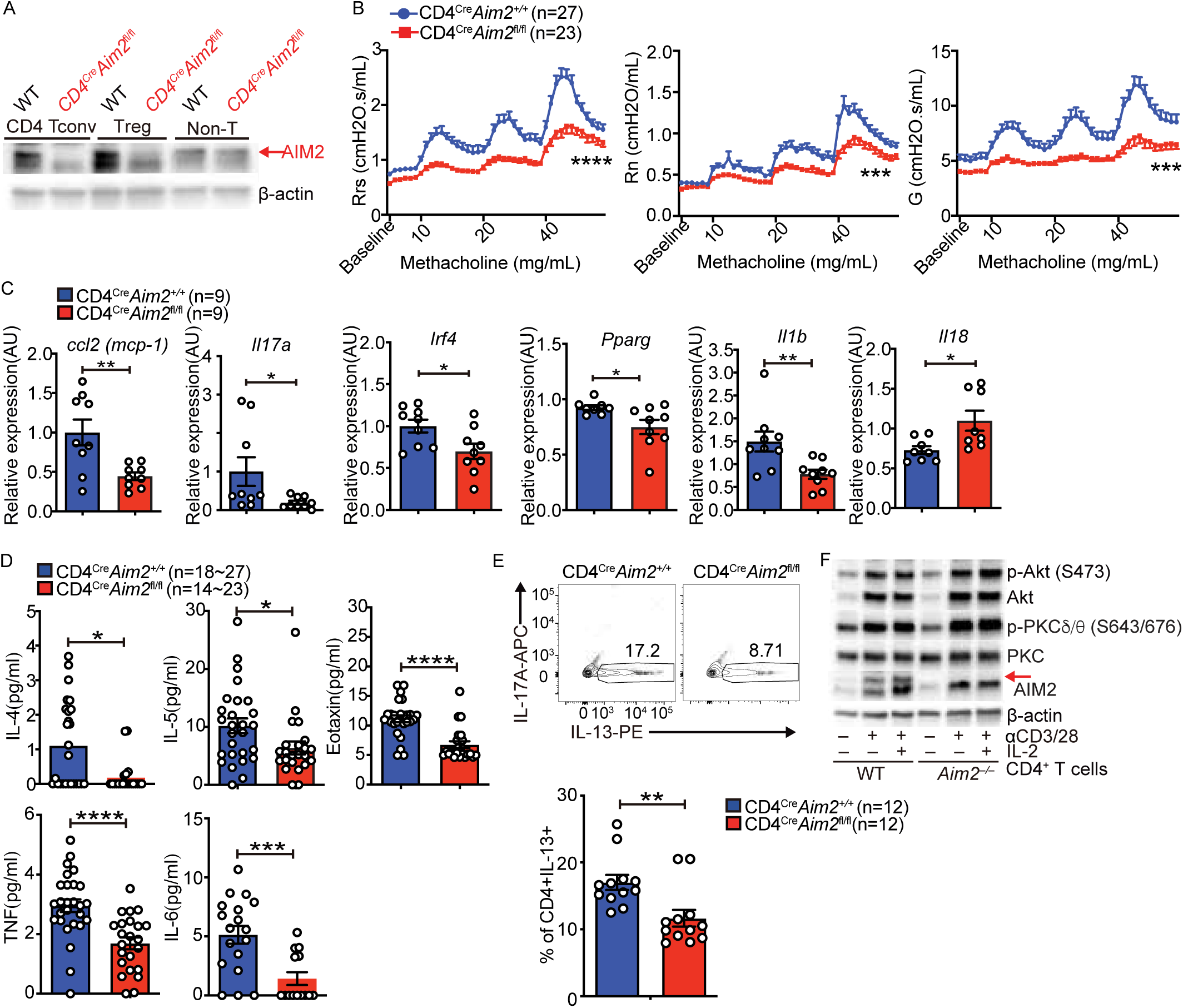
T cell-intrinsic but not myeloid-specific deletion of *Aim2* alleviates OVA-LPS-induced lung inflammation. (A) Immunoblot analysis of AIM2 protein in CD4 T^+^ cells, Tregs, and non-T cells from CD4^Cre^*Aim2^+/+^*and CD4^Cre^*Aim2*^fl/fl^mice. (B) Airway hyperresponsiveness (AHR) was measured upon administration of indicated methacholine doses, and presented by three parameters, including resistance of total airway (Rrs), proximal, large airway (Rn), and distal small airway (G) in mice. CD4^Cre^*Aim2^+/+^* n = 27, CD4^Cre^*Aim2*^fl/fl^n = 23. (C) Differential asthmatic-related inflammatory gene profiling was assessed in the lung. (D) Protein concentration of different cytokines in the BALF was measured by ELISA. (E) Expression of IL-13 in CD4⁺ T cells from CD4^Cre^*Aim2^+/+^* and CD4^Cre^*Aim2*^fl/fl^mice after 4 days of Th2 differentiation. (n = 6 mice/group; duplicate wells). (F) Immunoblot analysis of p-AKT(S473), AKT, p-PKCδ/θ (S643/676), PKCδ/θ, AIM2, and β-actin in wild-type and *Aim2*^−/−^ CD4^+^ T cells stimulated with anti-CD3/CD28 plus IL- 2 (40 U/ml) for 24 h. Data are representative of three independent experiments. Data shown represent mean values ± SEM. * p < 0.05; ** p < 0.01; *** p < 0.001; **** p < 0.0001. Student t- test or Two-way ANOVA with Sidak multiple comparison tests.

We next generated mice with a myeloid cell-specific deletion of *Aim2* by breeding LysMCre mice with a floxed *Aim2* allele. Myeloid-specific deletion of *Aim2* was confirmed by western blot (Fig. S3A). However, we observed no significant differences in AHR although there is a trend of reduction after administration of 20 mg/mL methacholine, and a modest but significant reduction in IL-5 production in the BALF of LysM^Cre^*Aim2* ^fl/fl^ mice, while the production of other asthma-related inflammatory cytokines including IL-1β and total antibody levels were unaffected (Fig. S3B-D). These results indicate that AIM2 in T cells is a critical driver of lung inflammation in OVA-LPS-induced allergic asthma model, while myeloid AIM2 has a minor role.

### AIM2 Expression is Elevated and Aggravates Lung Inflammation in the HDM-Induced Asthma Model

While OVA is the most widely used allergen to induce both acute and chronic experimental allergic asthma in mice, the use of an allergen that is relevant to human is important (*3, 11*). To further validate the role of AIM2 in regulating allergic asthma, we employed a house dust mite (HDM) model (Fig. 5A) as a second allergic asthma model. The HDM model stimulates both innate and adaptive immune responses and more accurately reflects the physiological characteristics of heterogeneous human asthma (*3, 11*). We observed increased *Aim2* expression in total lung homogenates from the HDM-induced group compared to the PBS control group (Fig. 5B). Similar to the phenotypes observed in the OVA-LPS model, *Aim2^−/−^* mice exhibited reduced AHR compared to WT mice in HDM-induced asthma model (Fig. 5C). Collectively, these results suggest that AIM2 regulates both OVA-LPS-induced asthma but also HDM-induced asthma.

**Fig. 5.**
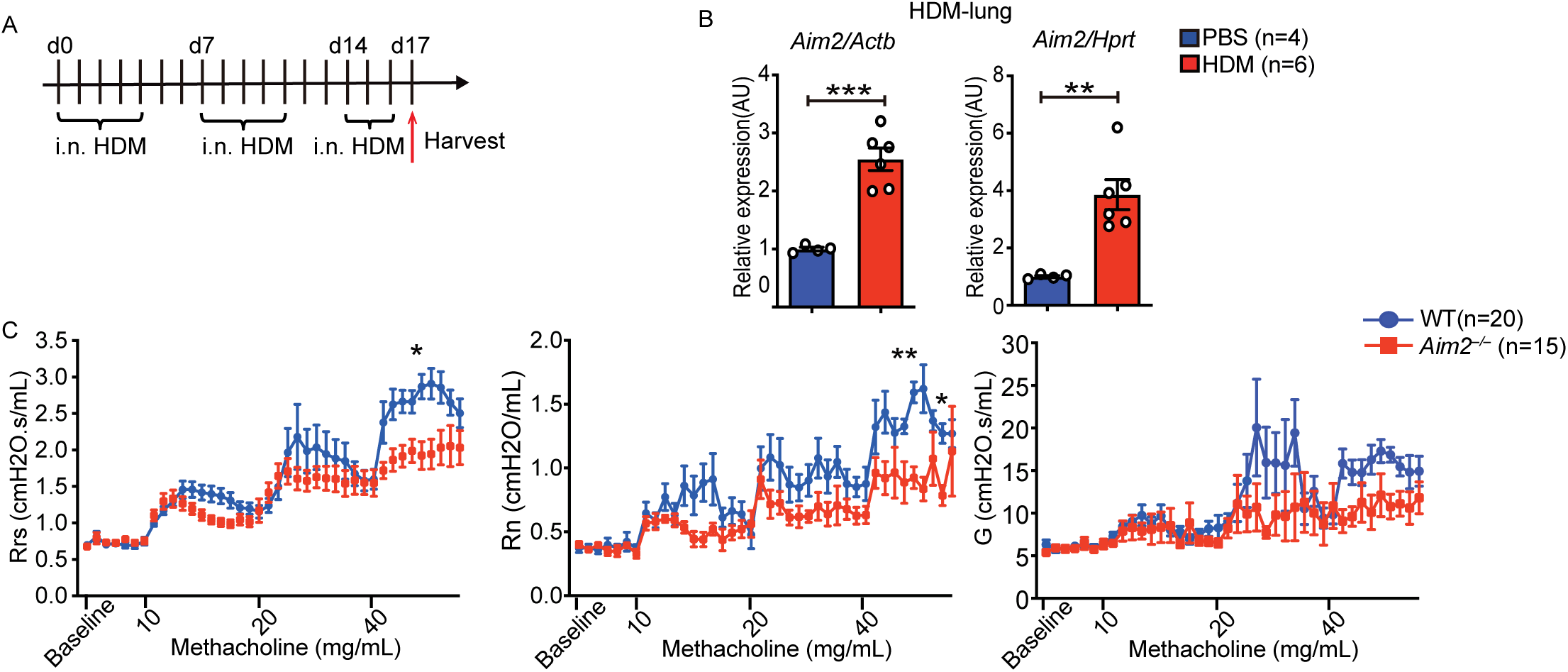
AIM2 exhibits increased expression and exacerbated lung inflammation in the house dust mite (HDM)-induced asthma model. *WT*, and *Aim2^−/−^* mice were continuously, intranasally sensitized to HDM (25 ng) for two and half weeks before harvest. (A) Schematic of HDM sensitizations, and harvest. (B) *Aim2* expression in the lung. (C) Airway hyperresponsiveness (AHR) was measured upon administration of indicated methacholine doses, and presented by three parameters, including resistance of total airway (Rrs), proximal, large airway (Rn), and distal small airway (G) in mice. *WT* n = 20, *Aim2^−/−^* n = 15. Data are representative of two independent experiments. Data shown represent mean values ± SEM. * p < 0.05; ** p < 0.01; *** p < 0.001. Student t-test or Two-way ANOVA with Sidak multiple comparison tests.

### T Cell-Intrinsic and Myeloid-Intrinsic Deletion of AIM2 Attenuates HDM-Induced allergic Asthma

Given the consistent reduction in AHR in *Aim2^−/−^* mice in HDM-induced asthma, we next investigated whether AIM2 influences the development of HDM-induced allergic asthma by regulating either myeloid cells or T cells. To determine whether AIM2 regulates the severity of HDM-induced allergic asthma in a T cell-intrinsic or myeloid cell-intrinsic manner, we applied the HDM-induced allergic asthma model to CD4^Cre^*Aim*2^fl/fl^, *Aim2*^fl/fl^FGCR26T(*43*), and LysM^Cre^*Aim*2^fl/fl^ mice to investigate AIM2’s role in cell-type-specific asthma regulation. The *Aim*2^fl/fl^FGCR26T mice has a Foxp3-specific deletion of *Aim2*, and hence *Aim2* is only deleted in T regulatory cells (Treg)(*43*). In CD4^Cre^*Aim*2^fl/fl^ mice, we consistently observed significantly reduced AHR (Fig. 6A). By contrast, there were no difference in AHR between *Aim2^+/+^*FGCR26T and *Aim*2^fl/fl^FGCR26T (Fig. 6B). We further found decreased production of IL-4 and IL-5 (Fig. 6C) but similar levels of Eotaxin, total IgE, IgG1, IgG2a production in CD4^Cre^*Aim*2^fl/fl^mice compared to CD4^Cre^*Aim2^+/+^*control mice (Fig. 6D). In LsyM^Cre^*Aim*2^fl/fl^ mice, we consistently observed reduced AHR (Fig. 6E), although no significant difference was noted in the levels of IL- 4, IL-13, and Eotaxin production between LysM^Cre^*Aim2^+/+^* and LysM^Cre^*Aim*2^fl/fl^ mice (Fig. 6F). Interestingly, we observed reduced serum total IgG1 production (Fig. 6G) and a decrease in IL- 13-producing CD4^+^ T cells and total alveolar macrophages (AMs) (Fig. 6H) in LysM^Cre^*Aim*2^fl/fl^mice compared to LysM^Cre^ *Aim2^+/+^*mice. These findings suggest that, in response to HDM stimulation, AIM2 differentially modulates immune responses through both myeloid cells and CD4^+^ T cells to promote airway hyperresponsiveness in allergic asthma.

**Fig. 6.**
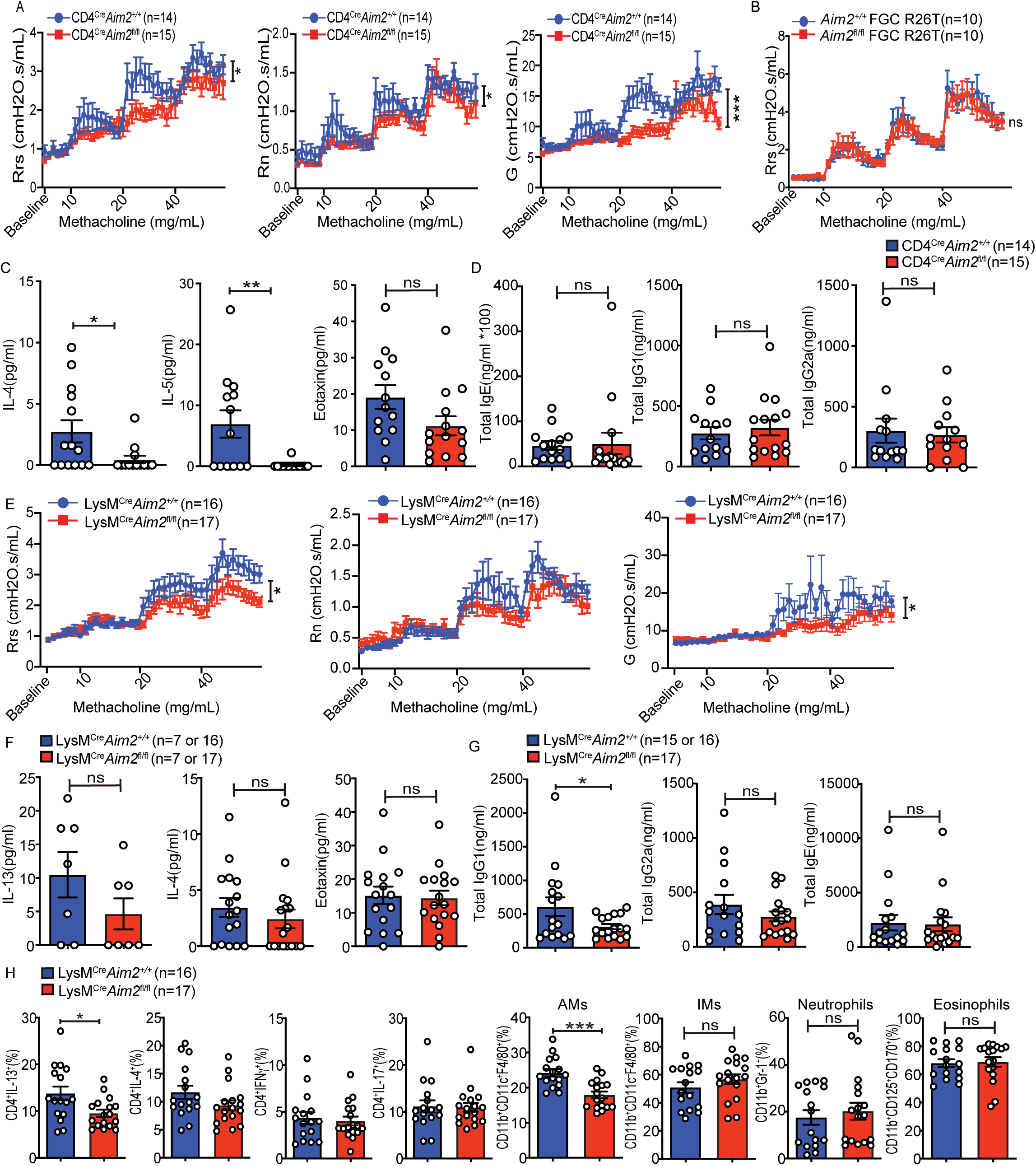
T cell-intrinsic and myeloid-intrinsic deletion of *Aim2* attenuate HDM-induced asthma. (A) Airway hyperresponsiveness (AHR) was measured in CD4^Cre^*Aim2^+/+^* and CD4^Cre^*Aim2*^fl/fl^mice upon administration of indicated methacholine doses, and presented by three parameters, including resistance of total airway (Rrs), proximal, large airway (Rn), and distal small airway (G) in mice. (B) Airway hyperresponsiveness (AHR) was measured in *Aim2^+/+^*FGC R26T and *Aim2fl/fl* FGC R26T mice upon administration of indicated methacholine doses and presented by resistance of total airway (Rrs). (C) Protein concentration of different cytokines in the BALF was measured by ELISA. (D) Serum IgG and IgE levels were determined by ELISA. CD4^Cre^*Aim2^+/+^*n = 14, CD4^Cre^*Aim2*^fl/fl^n = 15. Data are representative of two independent experiments. Data shown represent mean values ± SEM. * p < 0.05; ** p < 0.01; *** p < 0.001. Student t-test or Two-way ANOVA with Sidak multiple comparison tests. (E) Myeloid cell-intrinsic deletion of *Aim2* in the HDM-induced asthma. Airway hyperresponsiveness (AHR) was measured upon administration of indicated methacholine doses, and presented by three parameters, including resistance of total airway (Rrs), proximal, large airway (Rn), and distal small airway (G) in mice. (F) Protein concentration of different cytokines in the BALF was measured by ELISA. (G) Serum IgG and IgE levels were determined by ELISA. Data are representative of two independent experiments. (H) Flow cytometry analysis of innate immune cells and cytokine-producing CD4^+^ T cells in the BALF. LysM^Cre^*Aim2^+/+^* n = 16, LysM^Cre^*Aim2*^fl/fl^ n = 17. Data are representative of two independent experiments. Data shown represent mean values ± SEM. * p < 0.05; *** p < 0.001. Student t-test or Two-way ANOVA with Sidak multiple comparison tests.

## Discussion

Although the NLRP3 inflammasome has been extensively studied in asthma, its role has not always been consistent among studies. In this study, we investigated the role of AIM2 in allergic asthma using OVA-LPS and HDM-induced murine models, highlighting the immune cell-specific contributions of AIM2 to lung inflammation. Our findings indicate that AIM2 expression is elevated in both lung homogenates and BALF cells in these two asthma models compared to controls. In the OVA-LPS model, AIM2 exacerbates lung inflammation, with its deficiency leading to reduced airway hyperresponsiveness (AHR) and decreased levels of pro-inflammatory cytokines, including IL-4, CCL2, and TNF, accompanied by reduced Treg, and increased IFNγ^+^ CD4^+^ T cells. In the HDM-induced model, AIM2 deficiency similarly reduced AHR underscoring AIM2’s role in allergen-specific inflammatory regulation. We further show that AIM2 modulates immune cell populations in a cell-type-specific manner, and unexpectedly its expression in T cells had the most profound effects in both models, ultimately impacting asthma severity. By contrast, *Aim2* in myeloid cells showed a negligible role in the OVA-LPS model and a modest role in the HDM model.

Recent research shows that AIM2 expression is upregulated in severe pediatric asthma patients compared to healthy controls (*48*), while previous reports also indicate increased NLRP3 expression in the lungs of asthma patients and OVA-LPS-induced asthmatic mice compared to healthy controls (*22, 24, 67*). Our data indicate increased *AIM2* but not *NLRP3* expression in severe asthma. A similar pattern was observed when comparing neutrophilic asthma to paucigranulocytic asthma in human samples although NLRP3 showed a trend of increased expression in the former group of samples that did not reach a significant *p* value. This suggests that AIM2 may function in different cell types that contribute to the heterogeneity of asthma phenotypes and serves as a potential biomarker for severe asthma. The specific cell types in which AIM2 is activated has been assumed to be myeloid cells. However, in the OVA-LPS-induced asthma model, we observed reduced type 2 cytokine, reduced CD4^+^IL-13^+^ cells, reduced CD4^+^Foxp3^+^ regulatory T cells but increased IFNγ-producing but decreased IL-13-producing CD4^+^ T cells in *Aim2*-deficient mice. The reduced Treg is consistent with our previous findings of AIM2 promoting Treg cells. However, IL-17A-producing CD4^+^ T cells were unaffected in the whole body knockout strain, indicating that AIM2 may play a more prominent role in regulating Th1/Th2 responses. CD4-specific deletion of *Aim2* had a much greater impact on HDR lung inflammation than the whole body knockout, and resulted in reduced type 2 cytokines including BALF IL-4/IL-5 but also reduced CD4^+^IL-13^+^ T cells and reduced *Il17a* level, the last is consistent with its reported promotion of Th17 cells (*58*). By contrast, myeloid-specific deletion of *Aim2* had no impact on OVA-LPS induced asthma. In the HDM-induced asthma model, both T- and myeloid-specific AIM2 exacerbated diseases. CD4^+^ T cell-specific deletion of *Aim2* significantly alleviated AHR, lung inflammation, type 2 cytokines (IL4, IL5) and *Il17A,* consistent with its Th17 promoting phenotype differentiation (*58*). The myeloid-specific *Aim2* gene deletion also reduced AHR and lung inflammation, resulting in reduced IL-13 producing CD4 T cells and reduced total IgG1. It is interesting that AIM2 has been linked to the enhancement of T follicular (TFH) cells, which are key regulators of B cell activation and antibody production, in systemic lupus erythematosus (SLE) (*68*). Whether this is linked to reduced total IgG1 is unclear since our observation was made in myeloid-specific *Aim2* deletion mice. In total, these studies highlight the consistent promotion of type 2 inflammatory responses by *Aim2* in T cells, but also reveal that *Aim2* in myeloid cells may play a disease- and context-dependent role in asthma. AIM2 in myeloid cells may contribute to the initial activation of the immune response and the recruitment of inflammatory cells to the lungs.

In addition to its role in immune cells, the absence of AIM2 in human lung epithelial cells, which can form the first line of defense against pulmonary pathogens and allergens, has been associated with increased IFN-β production (*48*). Notably, AIM2 has also been implicated in influenza-induced lung inflammation, a major trigger of asthma exacerbations. AIM2 activation during viral infection may exacerbate asthma symptoms. The interplay between AIM2 and type I and III interferons (IFNs) during influenza infection may further complicate the inflammatory environment in asthma patients, especially those with concurrent viral infections. Moreover, we found the reduction of CCL2 level in OVA-LPS induced asthma in the absence of AIM2 while high level of CCL2 has been implicated in asthmatic sputum and supernatant from ASM, which plays a key role in airway hyperresponsiveness and remodeling in asthma (*51*). The reduction of CCL2 and α-SMA expression in AIM2-deficient lungs suggests that AIM2 may promote airway smooth muscle hypertrophy and contribute to airway remodeling. Additionally, *Aim2* is highly expression in mast cells (https://biogps.org), which are known to release a variety of inflammatory mediators, including histamine, and to infiltrate into and interact with ASM(*69*). Furthermore, NLRP3 has been shown to contribute to severe, steroid-resistant asthma, highlighting the importance of different inflammasomes in distinct types of asthma.

In summary, our data indicate that AIM2 is a key factor in AHR and inflammation in both the OVA-LPS and HDM-induced models. AIM2 plays an important role in T cells by promoting type II cytokines and T cells, but it also plays a role in myeloid cells in the HDM model. These results suggest that AIM2 inhibitors, such as andrographolide (***70***) or 4-sulfonic calixarenes (***71***) could represent novel therapeutic approaches particularly for patients with severe asthma. Additionally, modulating AIM2 activity in combination with other treatments may provide a promising strategy to enhance asthma control in severe cases.

## Materials and Methods

### Experimental Animals

Wild-type (C57BL/6), *Aim2*^−/−^, *Aim2*^fl/fl^ mice were generated as previously described (*43*). *Aim2*^fl/fl^ mice were then crossed with CD4Cre mice, to generate CD4 T cell-specific deletion of *Aim2* (CD4^Cre^*Aim2*^fl/fl^) and control mice (CD4^Cre^*Aim2*^+/+^). Additionally, *Aim2*^fl/fl^ mice were crossed with LysMCre mice, to generate myeloid cell-specific deletion of *Aim2* (LysM^Cre^*Aim2*^fl/fl^) and control mice (LysM^Cre^*Aim2*^+/+^). All mice were housed and bred under specific pathogen–free conditions with controlled temperature and humidity in the animal facility at the University of North Carolina at Chapel Hill. All sex- and age- matched (9–12 weeks) mouse experiments were approved by Institution Animal Care and Use Committee of the University of North Carolina.

### Mouse Models of Allergic Asthma

To induce airway inflammation, we used both OVA-LPS and HDM models. For the OVA-LPS model, mice were intranasally sensitized to total 26 μl volume for each mouse with 100 μg OVA (Grade V; Sigma-Aldrich, St. Louis, MO) plus 100 ng lipopolysaccharide (LPS) (LPS-EB from E. coli O111:B4, InvivoGen) as the adjuvant on days 0, 6, and challenged with 100 μg OVA on the day 13, 24 hours before the mice were sacrificed for harvest (Fig. 2A). For the HDM model, mice were sensitized intranasally with HDM extract (25 ng) (*Dermatophagoides pteronyssinus*, Greer laboratories, Lenoir, NC) in 25 μL PBS for two and half weeks (Fig. 5A). Mice were harvested 24 h following the last i.n. administration of OVA or HDM Ag in each model.

## Measurement of Lung Function by Small Animal Ventilator

Mice were anesthetized with 2.5% tribromoethanol (Avertin), tracheostomized, and paralyzed with Atracurium besylate. Mice were then mechanically ventilated with a computer controlled small animal ventilator (Scireq, Montreal, Canada) at 300 breaths per minute. Mice were challenged with either saline or 10, 20, and 40 mg/mL methacholine (Sigma, St. Louis) which were delivered by ultrasonic nebulizer (Scireq, Montreal, Canada) for 30 seconds. During aerosol exposure, ventilation rates were reduced to 200 breaths per minute and 0.15 cc/kg tidal volume. Ventilation rates were resumed to original values following aerosol challenge. Forced Oscillatory Mechanics (FOM) were utilized to evaluate airway reactivity every 10 seconds for 3 minutes following each Methacholine challenge. Our analysis focused on Newtonian resistance (Rn), which reflects the resistance in the central airways, Respiratory system resistance (Rrs), which measures the total respiratory system resistance, and tissue dampening (G), which reflects energy dissipation in the alveoli.

### Bronchoalveolar lavage fluid (BALF) collection and evaluation of airway inflammation

Mice were euthanized, and serum was collected from whole blood to measure total IgG1, IgG2a, and IgE levels. The lungs were lavaged two times with 1 ml of HBSS, and the collected bronchoalveolar lavage fluid (BALF) was centrifuged to separate the cellular components from the supernatants. Lungs were harvested and homogenized for RNA extraction and protein analysis. RNA levels in the lung homogenates were assessed using real-time PCR. Protein levels were analyzed from serum, cell-free BALF supernatants, and lung homogenates using ELISA (Biolegend; BD or R&D Biosystems) or Western blotting.

### RNA preparation and real-time PCR

The lung tissues and BALF cells were collected, and total RNA was prepared using TRIzol reagent (Invitrogen) per manufacturer’s instructions, and Zymo Direct-zol RNA Microprep (R2062), and then RNA samples were reverse-transcribed into cDNA with iScript^TM^ cDNA Synthesis Kit (Bio-Rad, cat.no. 1708891) for performing real-time PCR. The Taqman probes were purchased from Applied Biosystems (Thermo Fisher Scientific) or Integrated DNA Technologies (IDT), and quantitative PCR was performed on the ABI9700 real-time PCR system. Quantification of relative mRNA expression was normalized to the expression of *Actb* (Mm00261958) or *Hprt* (Mm01545399). Gene expression of mouse *Aim2* (Mm01295719), *Il1b* (Mm01336189), *Il18* (Mm00434225), *Il4* (Mm00445259), *Il5* (Mm00439646), *Il13* (Mm00434204), *Ifng* (Mm01168134), *Il17α* (Mm00439618), *Il33* (Mm00505403), *Ccl2* (Mm00441242), *Ifnb1* (Mm00439546), *Tnf* (Mm00443258), *Adam33* (Mm00459691), *Muc5ac* (Mm01276718), *Cxcl10* (Mm00445235), *Irf4* (Mm00516431), *Pparg* (Mm00440940), *Il6* (Mm00446190), *Il10* (Mm01288386), *Il12a* (Mm00434165), *Hmgb1* (Mm00849805), *Tbx21* (Mm00450960), *Rorc* (Mm01261022),and *Gata3* (Mm00484683) was analyzed using Taqman gene expression assays.

### Immunoblotting

The lung tissues were homogenized and lysed in Lysing Matrix D tubes (MP Biomedicals) containing RIPA buffer supplemented with complete proteinase inhibitor cocktail and PhoSTOP phosphatase inhibitors. The homogenization was performed using a FastPrep® homogenizer with the specified program settings. The procedure of western blot was followed the protocol as previously described (*43*). Briefly, the membranes were incubated overnight using the following primary antibodies (1:1000) from Cell Signaling Technology (CST): anti-GSDMD (ab209845), anti-p-Stat1 (Tyr 701) (cat.no. 9167), anti- Stat1 (cat.no. 9172), anti-p-Stat3 (Tyr 701) (cat.no. 9145), anti-Stat3 (cat.no. 9139), anti-AIM2 (cat.no. 63660), anti-cleaved caspase-3 (cat.no. 9664), anti-caspase-3 (cat.no. 9662), anti-cleaved caspase-7 (cat.no. 8438), anti-caspase-7 (cat.no. 12827), anti-p-H2A.X (Ser 139) (cat.no. 9718), anti-H2A.X (cat.no. 9361), anti-p-AKT (Ser473) (cat. no. 4060), anti-AKT (cat. no. 9272), anti-p-PKCδ/θ (S643/676) (cat. no. 9376), and anti-PKCθ (cat. no. 13643). The anti-alpha smooth muscle Actin (cat.no. ab7817) was from Abcam and anti-caspase-1 (p20, mouse; AG-20B-0042-C100) was from Adipogen. All primary antibodies were used at a 1:1,000 dilution in 1× TBS-T with 5% BSA. Membranes were washed in TBS-T and incubated with the following appropriate secondary antibodies from Jackson ImmunoResearch Laboratories: mouse anti-rabbit-HRP (211-032-171), donkey anti-rabbit HRP (711-035-152) or goat anti-mouse HRP (115-035-174). All secondary antibodies were used at a 1:10,000 dilution in 1× TBS-T with 5% BSA. Protein bands were visualized following exposure of the membranes to ECL substrate solution (ThermoFisher) and quantified by densitometric analysis using Image Lab software.

### The enzyme-linked immunosorbent assay (ELISA)

BALF cytokines and serum antibody production were analyzed by Enzyme-Linked Immunosorbent Assay (ELISA, MCS00, R&D Systems, Minneapolis, MN). ELISA kits were obtained from Biolegend for IL-4 (431104), IL-5 (431204), IL-13 (431104), IL-6 (431304), IL-1β (430642), TNF (430904), CCL-2/MCP-1 (432704), and IgE (431104); from R&D Systems for CXCL-10/IP-10 (DY466-05) and CCL-11/Eotaxin (DY420-05); and from Thermo Fisher Scientific (Invitrogen) for IL-18 (BMS618-3), IgG1 (88-50410-88), and IgG2a (88-50420-88). All assays were performed according to the manufacturers’ instructions.

### *In vitro* Th2 cell differentiation

CD4⁺ T cells were isolated from pooled peripheral lymph nodes and spleens of age- and sex-matched mice using CD4 (L3T4) microbeads (Miltenyi Biotec). Cells were cultured in RPMI 1640 supplemented with 10% FBS, 1% penicillin-streptomycin, and 2.6 μl/ml β-mercaptoethanol. Activation was performed with plate-bound anti-CD3 (2.5 μg/ml, 145-2C11, BioXCell) and soluble anti-CD28 (1 μg/ml, 37.51, BioXCell). For Th2 cell polarization, 20 ng/ml IL-4 (Biolegend), IL-2 (100 U/ml) and 20 μg/ml anti-IFNγ (XMG1.2, BioXcell) were added to the culture.

### Flow cytometry

BALF cells were collected on the day of harvest day after asthma induction. Fluorescence-conjugated antibodies for CD4 (GK1.5), CD8 (53-6.7), IFNγ (XMG1.2), IL-17A (TC11-18H10.1), IL-4 (11B11), IL-13 (W17010B), Foxp3 (MF-14), CD11b (M1/70), CD11c (N418), Gr-1(RB6-8C5), F4/80 (BM8), CD125 (DIH37), CD170 (S17007L) were purchased from Biolegend. The eBioscience™ Foxp3 / Transcription Factor Staining Buffer Set (00-5523-00) was obtained from Thermo Fisher Scientific. For intracellular cytokine staining, BALF cells were stimulated for 4 hours with 50 ng/ml of phorbol 12-myristate 13-acetate (PMA) and 1 mM ionomycin in the presence of brefeldin A. Stained cells were analyzed on an LSRFortessa station (BD Biosciences) using FACSDiva software. Intracellular cytokine staining was performed using a commercially available kit, following the manufacturer’s protocol (BD Biosciences). FACS data were analyzed with FlowJo software (TreeStar).

### Statistical analysis

Data analysis was performed and represented using Prism (GraphPad, San Diego). Statistical significance was assessed using a two-sided Student’s t-test or two-way ANOVA followed by Tukey’s post hoc test, as indicated in the figures. A p-value of less than 0.05 (95% confidence interval) was considered statistically significant. Asterisks in the figures represent p-values as follows: *p < 0.05, **p < 0.01, ***p < 0.001, and ****p < 0.0001. Sample sizes (n) are specified in the figure legends to indicate biologically independent replicates used for statistical analyses.

## Acknowledgements

We acknowledge funding support from the NIH (AI029564, AI158314 to J.P.-Y.T.; AI160774, AI123193 to Y.Y.W and W.C.; HL150541 to L.B.A), and the National Multiple Sclerosis Society (NMSS FG 1968-A-1 to M.H.). We thank Dr. Dale Cowley and the UNC animal core for generating *Aim2*^fl/fl^ mice, Dr. Willie June Brickey for managing the *Aim2*^fl/fl^ mice, and Dr. Xiaoqing Hu for immunoblotting assistance.

## Author Contributions

W.C.C and J.P.-Y.T designed research; W.C.C, M.H., E.H.G., C.M.J and K.L. performed research; W.C.C., A.L.B., S.L.T, Y.Y.W, contributed new reagents/analytic tools; W.C.C, M.H., C.M.J, K.L. and J.A.W. analyzed data; and W.C.C and J.P.-Y.T wrote the paper.

## Supplementary Information

**Figure S1.**
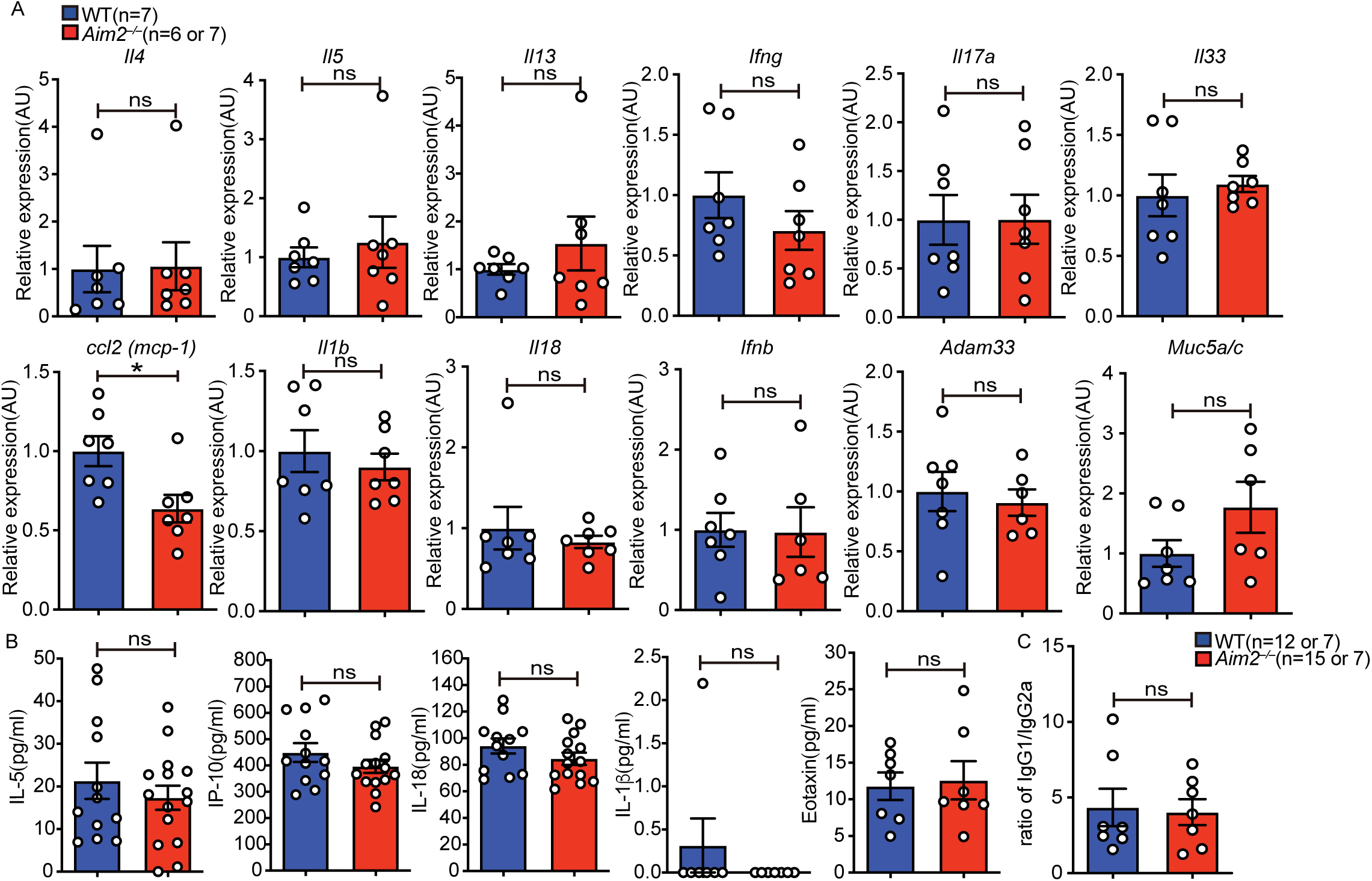
Comparison of differential asthmatic related gene profile, cytokine expression, and antibody production WT and *Aim2^−/−^* mice were. (A) Real-time PCR was performed to exam gene expression in lung. (B) Proinflammatory cytokines and chemokines and (C) the ratio of serum IgG1 and IgG2a titers were measured by ELISA. Data shown represent mean values ± SEM. ns: p >0.05; not significant by Student t-test.

**Figure S2.**
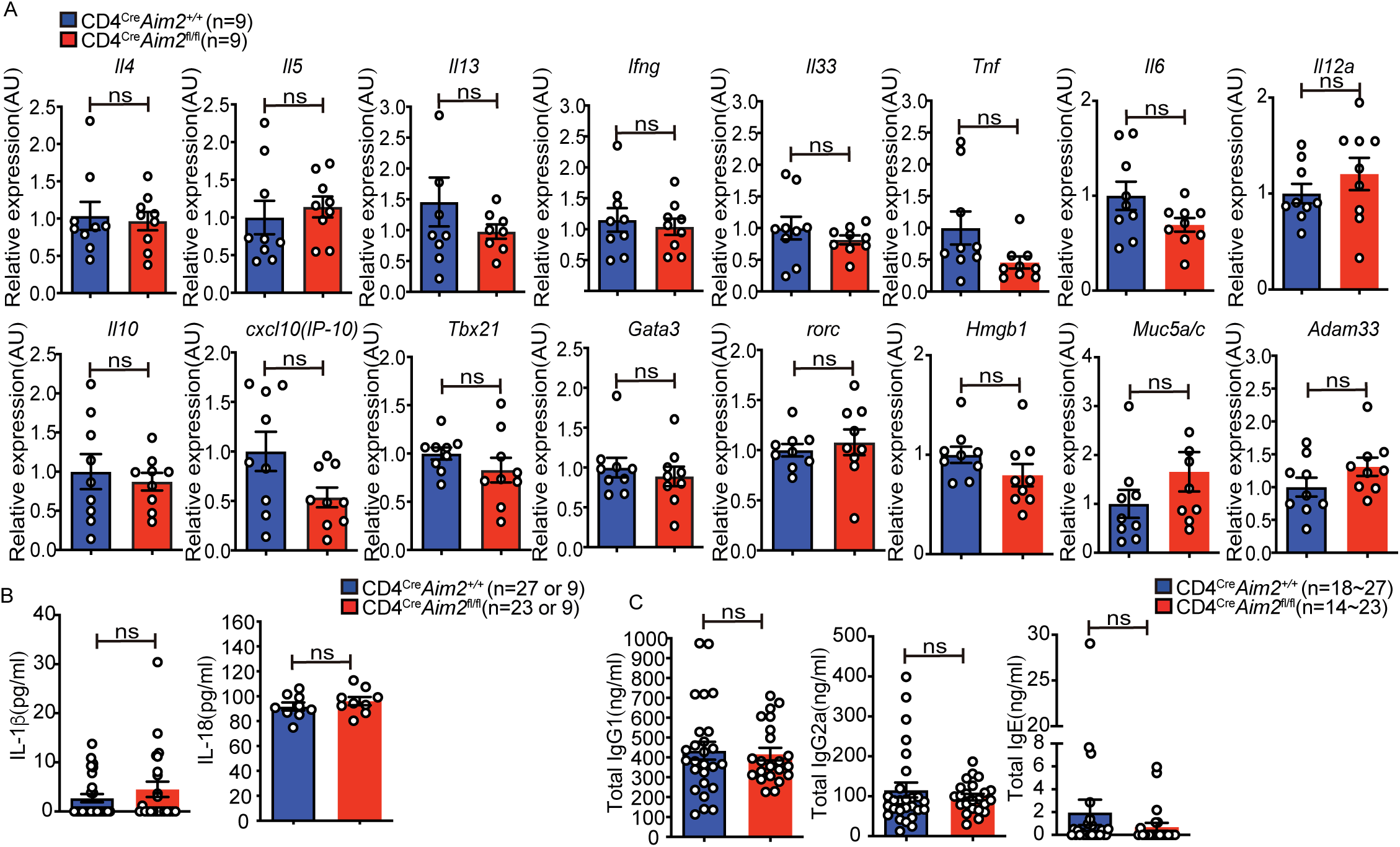
Comparison of differential asthmatic related gene profile, cytokine expression, and antibody production between CD4^Cre^*Aim2^+/+^* and CD4^Cre^*Aim2*^fl/fl^mice. (A) Real-time PCR was performed to exam gene expression in the lung. (B) The protein levels of IL-1β and IL-18 and (C) the concentration of serum IgG1, IgG2a and IgE were measured by ELISA. CD4^Cre^*Aim2^+/+^* n = 9 or 18∼27, CD4^Cre^*Aim2*^fl/fl^ n = 9 or 14∼23. Data shown represent mean values ± SEM. ns: p >0.05; not significant by Student t-test.

**Figure S3.**
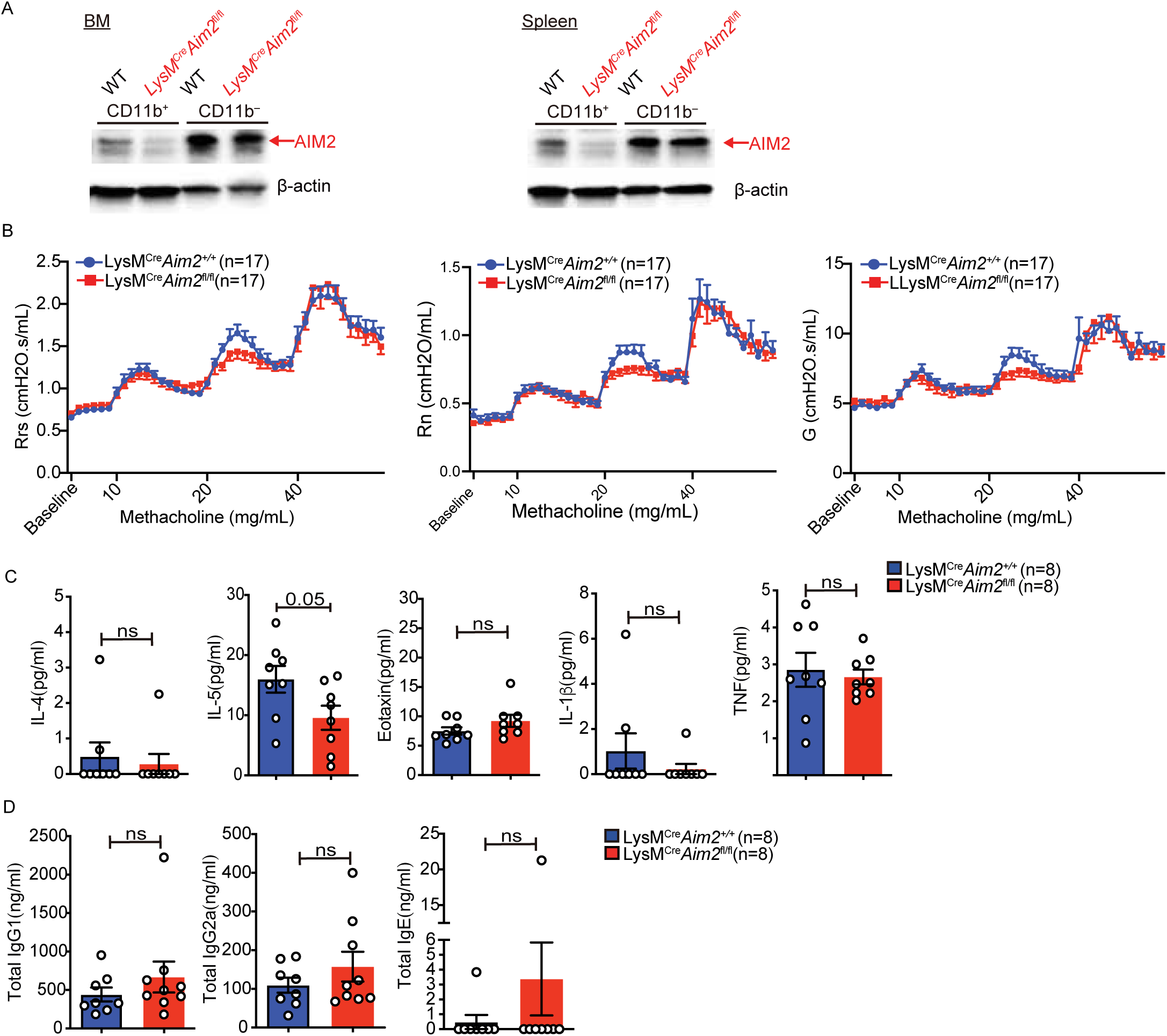
Lack of AIM2 in myeloid cells has no effects in the OVA-LPS model. (A) Immunoblot analysis of AIM2 protein in CD11b^+^ and CD11b^−^ cells purified by CD11b beads from total bone marrow (BM) or splenocytes (SPN) in LysM^Cre^*Aim2^+/+^*and LysM^Cre^*Aim2*^fl/fl^ mice. (B) Airway hyperresponsiveness (AHR) was measured upon administration of indicated methacholine doses, and presented by three parameters, including resistance of total airway (Rrs), proximal, large airway (Rn), and distal small airway (G) in mice. LysM^Cre^*Aim2^+/+^* n = 17, LysM^Cre^*Aim2*^fl/fl^ n =17. (C) Protein concentration of different cytokines in the BALF was measured by ELISA. (D) Serum IgG and IgE levels were determined by ELISA. LysM^Cre^*Aim2^+/+^* n = 8, LysM^Cre^*Aim2*^fl/fl^ n = 8. Data are representative of two independent experiments. Data shown represents mean values ± SEM. p = 0.05. ns: p >0.05; not significant. Student t-test or Two-way ANOVA with Sidak multiple comparison tests.

